# Deep learning-based 3D spatial transcriptomics with X-Pression

**DOI:** 10.1101/2025.03.21.644627

**Authors:** Demeter Túrós, Lollija Gladiseva, Marius Botos, Chang He, G. Tuba Barut, Inês Berenguer Veiga, Nadine Ebert, Anne Bonnin, Astrid Chanfon, Llorenç Grau-Roma, Alberto Valdeolivas, Sven Rottenberg, Volker Thiel, Etori Aguiar Moreira

## Abstract

Spatial transcriptomics technologies currently lack scalable and cost-effective options to profile tissues in three dimensions. Technological advances in microcomputed tomography enabled non-destructive volumetric imaging of tissue blocks with sub-micron resolution at a centimetre scale. Here, we present X-Pression, a deep convolutional neural network-based framework designed to reconstruct 3D expression signatures of cellular niches from volumetric microcomputed tomography data. By training on a singular 2D section of a paired spatial transcriptomics experiment, X-Pression achieves high accuracy and is capable of generalising to out-of-sample examples. We utilised X-Pression to demonstrate the benefit of 3D examination of tissues on a paired SARS-CoV-2 vaccine efficacy spatial transcriptomics and microcomputed tomography cohort of a recently developed live attenuated SARS-CoV-2 vaccine. By applying X-Pression to the entire mouse lung, we visualised the sites of viral replication at the organ level and the simultaneous collapse of small alveoli in their vicinity. In addition, we assessed the immunological response following vaccination and virus challenge infection. X-Pression offers a valuable and cost-effective addition to infer expression signatures without the need for consecutive 2D sectioning and reconstruction, providing new insights into transcriptomic profiles in three dimensions.

## Introduction

Spatial transcriptomics (ST) technologies have had a massive impact on life sciences in recent years, ranging from new findings in developmental biology to cancer research with discoveries directly linked to patient outcomes [1–3]. *In situ* profiling technologies enable us to examine cells and their organisation in the tissue space, revealing new patterns and spatial interactions [4–6].

Despite the rapid development of numerous ST technologies, including *in situ* sequencing and spatial barcoding approaches, most of these platforms struggle with scalability due to proprietary components and high costs [7]. Consequently, together with the histopathology standards that involve working with thin two-dimensional tissue sections, these platforms tend to overlook that tissues are threedimensional structures.

Novel methodologies and computational approaches in the field of ST are exploring more cost-effective open-source platforms allowing serial sectioning and 3D reconstruction [8–12]. Although these approaches led to significant discoveries by examining the 3D structure, they still require multiple thin sections to be processed and reconstructed, which in turn restricts their scalability and results in the destruction of the valuable original tissue samples. New generations of imaging mass cytometry [13] and MERFISH [14] are being developed to effectively analyse thicker (200-300 µm) tissue samples. While these methods offer crucial insights into 3D spatial biology, they present several technological drawbacks, such as limited tissue penetration of antibodies/oligos [15] and diminished signal intensity [14, 16].

The field of computational pathology has been adapting to emerging 3D imaging technologies that enable the capture of volumetric tissue data in a non-destructive manner [17, 18]. Modalities such as light-sheet microscopy [19], multiphoton microscopy, optical coherence tomography [20], photoacoustic microscopy [21], and microcomputed tomography (micro-CT) have been shown to enhance standard pathological workflows. Among these, micro-CT is capable of imaging large tissue samples (centimetre scale) at sub-micrometre resolution directly on the paraffin blocks [22, 23]. This technology has also demonstrated success in resolving the 3D microstructure of solid tissues [24] and consistently produced quantitative features without disrupting histological workflows [25].

Deep learning-based computational tools have shown great promise in linking histology to gene expression. Various methods, including as ST-Net [26], XFuse [27], and iStar [28] have been developed to predict gene expression from haematoxylin and eosin (H&E) tissue sections or to enhance the resolution of traditional ST platforms by imputing gene expression from histology.

Building on these approaches, we introduce X-Pression, a novel deep learning-based framework designed to reconstruct gene expression programs in 3D from micro-CT data. We demonstrate its effectiveness using a cohort of SARS-CoV-2 infected mouse lungs profiled with both 10x Visium and micro-CT. X-Pression reliably infers expression programs related to viral infection and replication using a single ST sample for training. Additionally, we show how investigating 3D spatial domains can provide valuable insights into SARS-CoV-2 infection, disease progression, and the efficacy of various vaccines. Our approach also allows for accurate prediction of expression profiles in volumes that lack paired ST data from the same cohort. X-Pression offers a scalable, costeffective alternative for exploring the 3D transcriptional landscape of tissues, paving the way to new discoveries in both basic and applied research.

## Results

### Model overview

To enable scalable and cost-effective inference of 3D gene expression programs from volumetric imaging data, we developed X-Pression. This framework builds on a deep convolutional neural network (CNN) backbone that takes 3D patches from the micro-CT volume to predict gene expression programs. First, we obtained tissue sections from paraffin blocks containing fixed tissues for ST with 10x Visium, while the remaining blocks were used for micro-CT scanning (**Fig.1a**). Next, we preprocessed the micro-CT volumes by segmenting them with a U-net architecture trained on axial tomogram slices annotated by veterinary pathologists. After co-registering the two modalities with landmark-based affine transformations, X-Pression extracted 3D volumetric patches (128×128×21 pixels; 204.8×204.8×33.6 µm) centred around ST capture spots from the micro-CT volume. By extending the patch size over the boundaries of the capture spots (diameter = 55 µm), we incorporated additional spatial context that matches the average distance of paracrine signalling molecules (200-300 µm) capturing the potential influence of the surrounding tissue [29, 30]. These patches are subsequently used to train a supervised CNN to infer gene expression programs associated with specific cellular niches (*i*.*e*., tissue compartments) calculated by Chrysalis [31] (**Fig.1b**). In the CNN, 3D convolutional layers perform feature extraction, followed by a regressor head with a final output layer that employs a softmax activation function. Since the expression programs calculated with Chrysalis are represented as a convex combination summing to one, the softmax outputs are directly used as their predicted values for each capture spot. Hence, by leveraging deep learning and spatial context beyond individual capture spots, X-Pression enables a robust, high-resolution reconstruction of gene expression profiles in 3D.

**Fig. 1.**
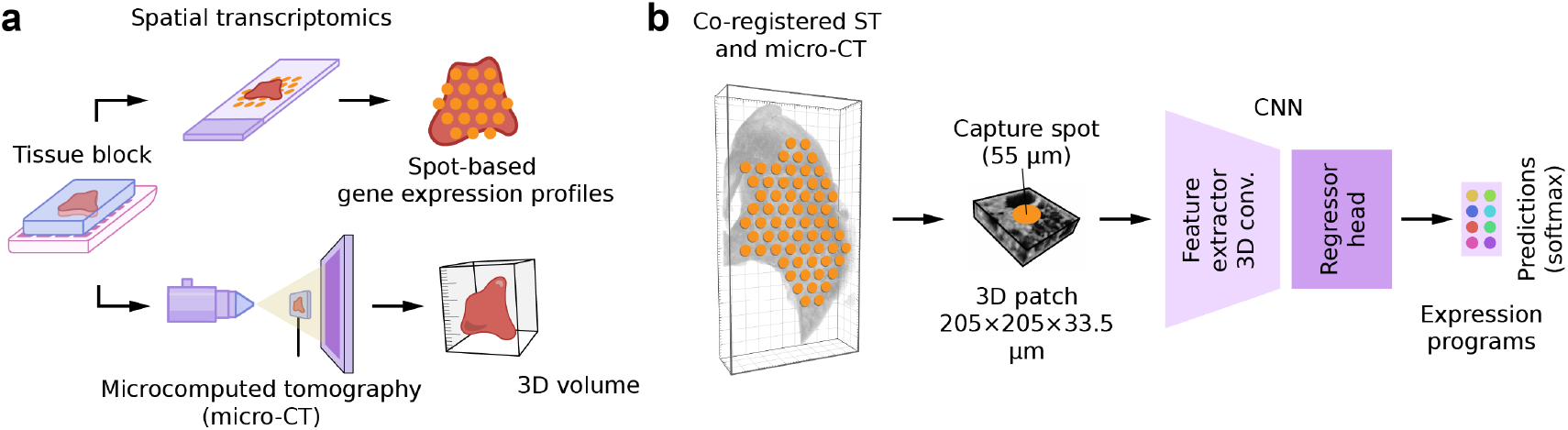
X-Pression overview. **a**, X-Pression framework leverages a formalin-fixed paraffin-embedded (FFPE) tissue block. A single tissue section is taken from the surface of a paraffin block and subjected to ST (10x Visium), while the rest of the block is imaged using micro-CT. This approach produces paired samples of spot-based ST data and micro-CT volumes of the whole tissue blocks with a resolution of 1.625 µm. **b**, After co-registration, X-Pression takes 3D image tiles from the micro-CT volume centred around each capture spot for training. The CNN model of X-Pression is composed of a feature extractor containing four blocks of 3D convolutional and max-pooling layers, followed by a regressor head with a softmax activation function to predict gene expression programs.

### X-Pression accurately infers gene expression programs from micro-CT data

To assess the accuracy and applicability of X-Pression, we applied it to an ST and micro-CT dataset from a severe acute respiratory syndrome coronavirus 2 (SARS-CoV-2) vaccine efficacy study. We analysed samples from animals immunised with either a genome-modified live-attenuated virus, including “one-to-stop” codons (OTS206) [32], or an mRNA vaccine. Following SARS-CoV-2 Delta (B.1.617.2) variant challenge, we compared their elicited immune responses in 3D.

First, we evaluated X-Pression using a single-pair approach on a specimen (L2210926) from the right lung of an mRNA vaccine-immunised mouse, collected 5 days post-challenge (5 dpc) with the SARS-CoV-2 Delta variant. Assessing the ST data with Chrysalis, we identified eight distinct gene expression programs that represent functionally and spatially distinct anatomical regions associated with cellular niches. These expression programs were used as training targets for X-Pression **(Fig.2a)**. We characterised these key expression programs by analysing their most representative and distinctly expressed genes **(Fig.2b-c)**, and by correlating them with various cell types via deconvolution relying on our single-cell reference data, histopathological annotations, and functional characterisation **(Supplementary Note 1-2)**. Further analysis conducted on the ST datasets (spatiotemporal assessment of the expression programs and pathway activity inference) of mouse lungs infected with either wild-type (Wuhan strain) SARS-CoV-2 and OTS206, or challenged with the Delta variant post-immunisation with an mRNA vaccine and OTS206 are described in **Supplementary Note 2**.

**Fig. 2.**
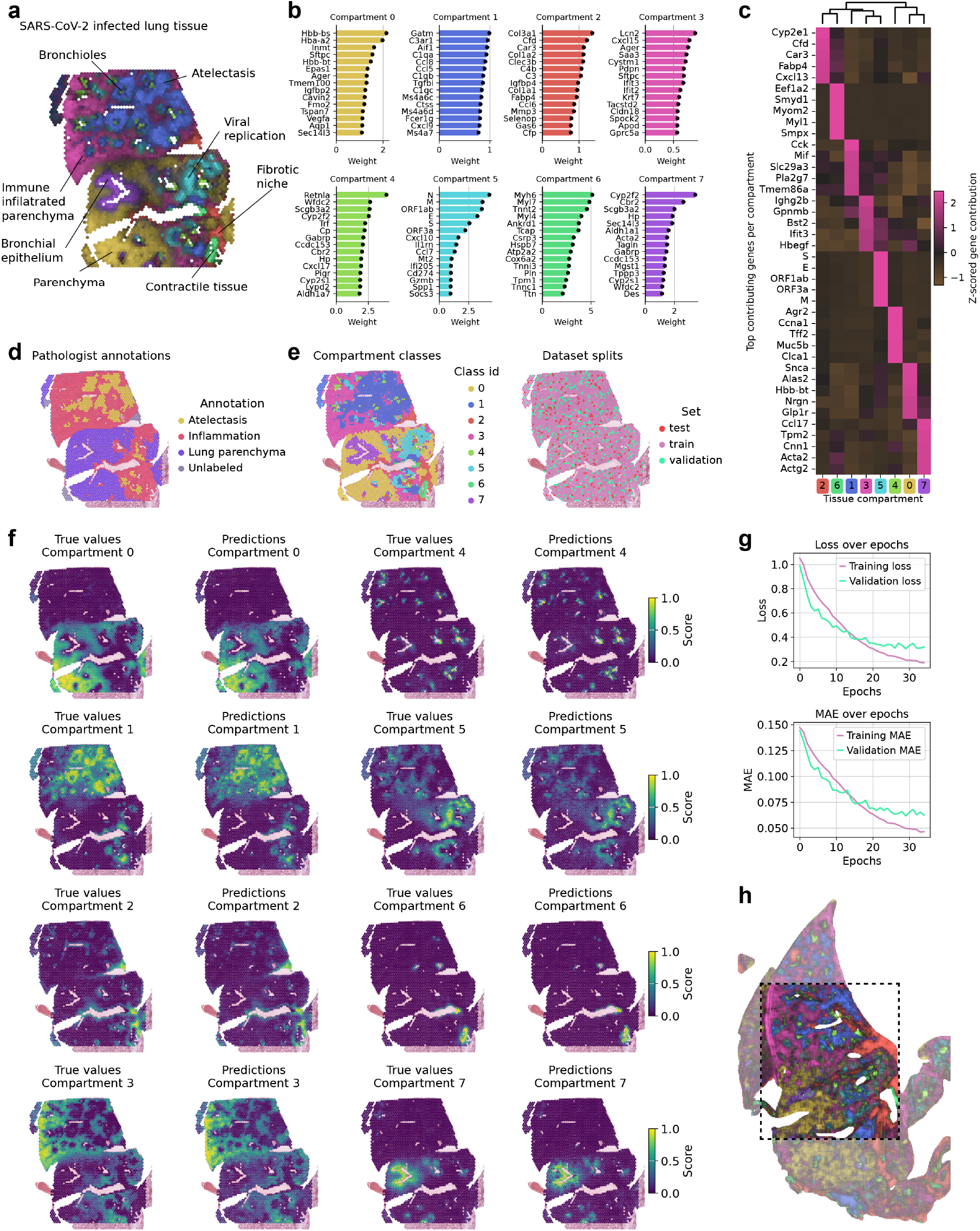
X-Pression infers gene expression programs on the organ level using a single ST section. Data originated from a specimen (L2210926) obtained from the right lung of an mRNA-vaccinated mouse five days after challenge with the SARS-CoV-2 Delta variant. **a**, Maximum intensity projection (MIP) of the gene expression programs. **b**, Highest-weighted genes in each expression program. **c**, Heatmap showing the Z-scored contribution of genes to the expression programs. **d**, Pathologist annotations based on the H&E image. **e**, Capture spots classified based on the highest gene expression program for stratified sampling (left) and spots belonging to training, test, and validation sets (right). **f**, True values for each gene expression signature per capture spot next to the values predicted by X-Pression. **g**, Changes in loss and MAE during the training process. **h**, MIP of the predictions on an axial section of the whole FFPE block. The dashed-lined rectangle corresponds to the position and area of the tissue section measured with ST.

The characterisation of the eight different gene expression programs revealed distinct patterns associated with lung architecture and cellular processes **(Fig.2a-b)**. The expression program linked to lung parenchyma (0) is characterised by genes primarily expressed in pulmonary alveolar cells. Additionally, parenchyma infiltrated with resident immune cells (3) was identified. Moreover, distinct expression signatures suggest the presence of a fibrotic and immune-modulated microenvironment (2), bronchioles (4), larger airways featuring contractile tissue (6), and bronchial epithelium (7). The signature of viral replication sites (5) was defined based on the expression of SARS-CoV-2 *N, M, ORF1, E*, and *S* genes. Additionally, the presence of immune markers such as *Cxcl10, Ccl7, Ifi205, Socs3*, and *Il1rn* indicated a robust antiviral response driven by interferons in the replication sites. The expression of *Cd274 (PD-L1)* and Gzmb implied the involvement of immune regulatory and activation processes, possibly through cytotoxic T cells or NK cells, alongside tissue remodelling and an oxidative stress response indicated by Spp1 and Mt2 expression. This was further confirmed by the upregulation of JAK-STAT and NfkB pathways **(Supplementary Note 2)**. An additional expression program, termed inflammation-induced atelectasis (IIA) (1), corresponding to areas of severe immune infiltration, which expanded the alveolar septa, occluded the alveolar lumina, and led to associated alveolar collapse was further identified. Genes associated with this signature are indicative of the presence of macrophages (Ctl8, Ctss), neutrophils (Aif1, Ccl5) and complement activation (C1 genes). In further correlation with TGF-β, these areas are consistent with severe histiocytic and neutrophilic inflammation resulting in airway remodelling and alveolar collapse. These major expression programs were in accordance with annotations based on histopathological analysis (**Fig.2d**).

After characterising the gene expression programs in the tissue sample, we trained X-Pression to infer these programs from the micro-CT data. Our model training strategy involved first extracting the capture spot-3D tile pairs that were subsequently classified for stratified sampling and data augmentation. Classes were defined based on the highest compartment score for each capture spot and split to train, test, and validation sets (**Fig.2e**). We evaluated the performance of X-Pression by calculating the coefficient of determination (*R*^2^) for these sets (**Table 1**). Overall, the total average *R*^2^ was 0.860, from which 0.643 was achieved on the validation and 0.583 on the test set. In the test set, predictions for multiple expression programs exceeded 0.7. Visualising the inferred values in the tissue space shows robust spatial patterns with high similarity to the ST data (**Fig.2f**). Training strategy involved monitoring the mean absolute error (MAE) in the validation set, which was concluded if no further improvements were measured in the last five epochs (**Fig.2g**).

**Table 1.** *R*^2^ for different splits.

| Comp. | Total $R^2$ | Train. | Val. | Test |
| --- | --- | --- | --- | --- |
| 0 | 0.9122 | 0.9338 | 0.7841 | 0.8183 |
| 1 | 0.9152 | 0.9348 | 0.8175 | 0.7502 |
| 2 | 0.8260 | 0.9086 | 0.5570 | 0.5868 |
| 3 | 0.8461 | 0.8894 | 0.6235 | 0.5907 |
| 4 | 0.7885 | 0.9281 | 0.3693 | 0.0616 |
| 5 | 0.8224 | 0.9023 | 0.5706 | 0.5486 |
| 6 | 0.8813 | 0.9377 | 0.7541 | 0.6291 |
| 7 | 0.8882 | 0.9597 | 0.6676 | 0.6811 |

To comprehensively map gene expression programs, we extracted 3D tiles from the micro-CT volume using a sliding window with a stride of 16 pixels (26 µm). This approach reduces aliasing artefacts and edge effects, resulting in capturing finer variations in gene expression. Reconstruction of the predicted signatures allowed us to visualise them on the whole organ level that reaches beyond the boundaries of the ST capture area (**Fig.2h**). These results highlight the strong predictive performance of X-Pression, with inferred gene expression programs that align closely with ST data. Our approach enables the mapping of spatial gene expression patterns, offering a robust framework for studying complex tissue environments.

### X-Pression reveals 3D infection patterns during SARS-CoV-2 infection

Next, we reconstructed the inferred gene expression programs in 3D, revealing complex spatial features across the lung tissue. Examination of an axial cross-section revealed the widespread presence of the IIA signature (1) in all major lobes of the right lung (**Fig.3a**). These structures were often surrounded by active viral replication sites, as indicated by the predictions of compartment 5, which is defined by SARS-CoV-2 genes and cytokine expression. Replication sites were also observed further away from atelectatic immune-infiltrated areas, indicating a more complex temporal relationship between active viral replication and lung collapse. In addition to the prominent virus-associated compartments, we observed strong signals of immune cell-infiltrated parenchyma (3) in the caudal lobe. Further examination of multiple axial sections revealed prominent viral replication sites and IIA throughout the entire lung (**Fig.3b**), with more foci of IIA in the centre of active viral replication. Cross-sectional and parasagittal volume slices (**Fig.3c-d**) further demonstrated this.

**Fig. 3.**
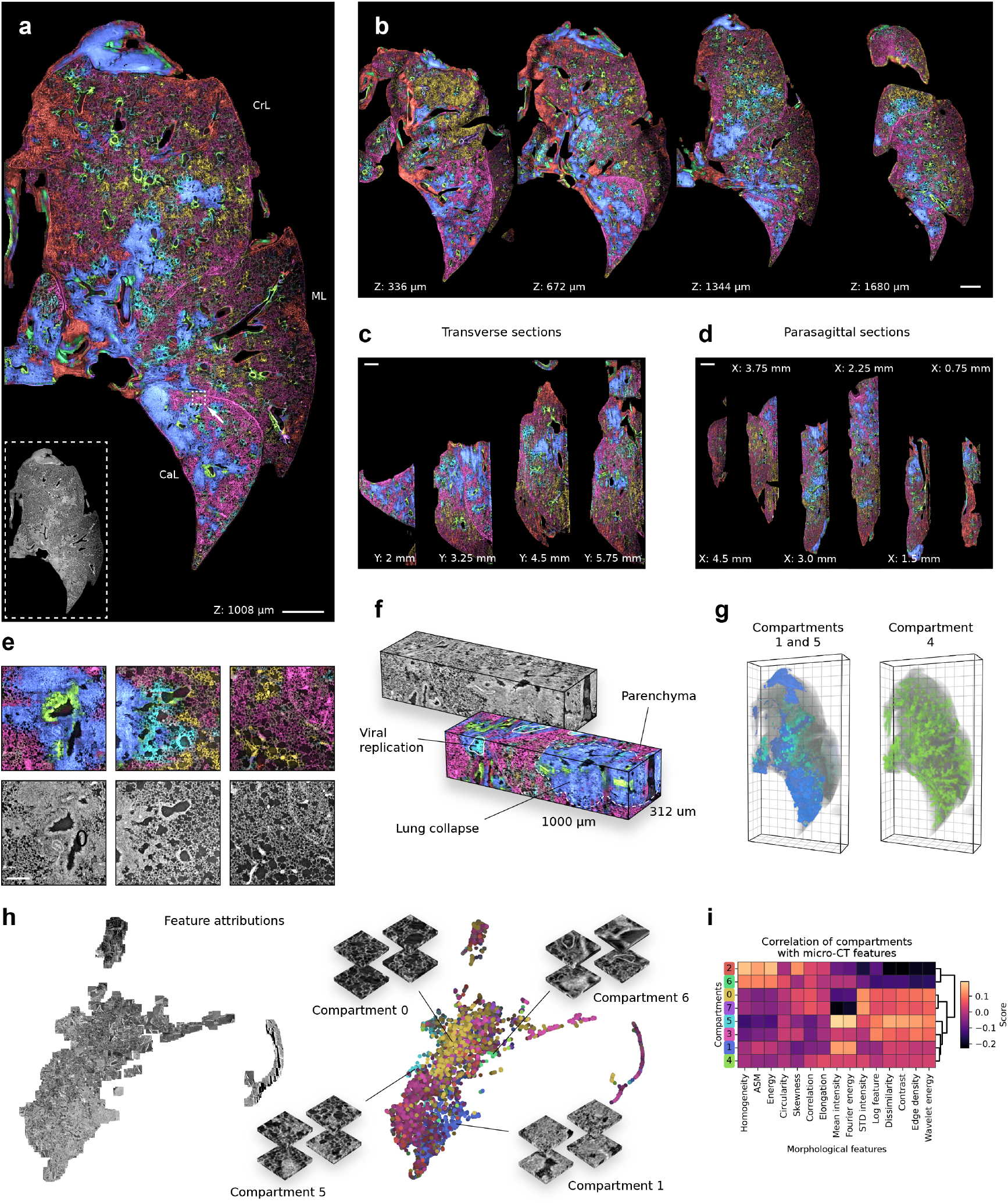
Gene expression signatures revealed with X-Pression in 3D during SARS-CoV-2 infection. **a**, Axial section of SARS-CoV-2-infected mouse lung (L2210926, mRNA vaccine, 5dpc) with MIP colour overlays showing the gene expression signatures inferred by X-Pression. The original tomogram section is shown in the bottom left inset. Dashed-lined square with a white arrow: sampling site for panel f. Scale bar: 1000 µm. CrL: cranial lobe, ML: medial lobe, CaL: caudal lobe. **b**, Serial axial sections with colour overlays showing gene expression changes across the volume. Scale bar: 1000 µm. **c**, Serial transverse sections with colour overlays showing gene expression changes across the volume. Scale bar: 1000 µm. **d**, Serial parasagittal sections with colour overlays showing gene expression changes across the volume. Scale bar: 1000 µm. **e**, Demonstrative high-resolution ROIs extracted from the axial section shown in panel a with colour overlays showing the gene expression signatures and the original tomogram. Scale bar: 200 µm. **f**, Cuboid sampled from the lung at the location indicated in panel a, where the drawn square corresponds to its base. **g**, 3D whole organ renders of expression programs associated with IIA (1) and viral replication (5) (left) and bronchioles (4) (right). Grid spacing: 500 µm. **h**, UMAP plots of IG attributions. On the left, individual data points are represented by their corresponding middle slices from the extracted 3D patches. On the right, points are coloured according to their predicted compartment scores. **i**, Heatmap of correlation matrix displaying the Pearson correlation coefficients between micro-CT morphological features and predicted tissue compartments.

We then focused on examining the microstructure in more detail (**Fig.3e**). On 500 µm wide square-shaped region-of-interests (ROIs), X-Pression accurately delineated areas of IIA, which were primarily visible on the tomogram as consolidated structures of high radiodensity. Additionally, viral replication sites were distinguished from the surrounding parenchyma. Contrary to normal alveolar structures, these areas correlated with distorted alveolar spaces demarcated by irregularly thickened alveolar septa corresponding to the interstitial immune cell infiltration in histology.

To demonstrate the flexibility of our approach, we extracted a square prism with a height of 1000 µm from the volume (**Fig.3f**). X-Pression revealed intricate gene expression changes across the prism, further exemplifying the method’s capacity to visualise complex spatial relationships between viral replication, immune cell infiltration, and tissue architecture.

Building on these analyses, we generated whole-organ 3D renders of the inferred expression programs (**Fig.3g**). The dual 3D render of lung collapse and viral replication revealed a prominent enrichment of the IIA-associated signature in the caudal lobe. It also highlighted the overall pattern of IIA following the major airways at the centre of active viral infection. Additionally, we created a separate render of the bronchus-associated expression signature, which effectively visualised the branching pattern of the airways.

Furthermore, we computed feature attributions using Integrated Gradients (IG) (**Fig.3h**). The attributions were then projected into a 2D UMAP space to visualise the relationship between tissue structure and model predictions. The resulting UMAP visualisation revealed distinct clusters that correspond to patches with varying textures and intensities, which are associated with different expression programs.

Finally, we computed Pearson correlation coefficients between micro-CT-derived morphological features and the predicted compartment values using patches from the entire micro-CT volume. Our analysis revealed that different expression signatures identified by the model align with specific morphological patterns (**Fig.3i**). Although the correlations were not pronounced, we observed that different compartments showed distinct morphological feature compositions. For example, compartment 7, which corresponds to a bronchial epithelium, displayed a negative correlation with both mean intensity and Fourier energy. This pattern is consistent with the structural properties of airways in micro-CT, which typically appear as low-intensity regions due to their thinner structure and reduced X-ray attenuation. In contrast, compartment 1, representing regions affected by IIA, showed a positive correlation with both mean intensity and Fourier energy, alongside more pronounced negative correlations with skewness, Laplacian of Gaussian (LoG), and standard deviation (STD) of intensity. IIA-affected regions appear denser and more homogeneous on micro-CT scans. This is consistent with the positive correlation between mean intensity and Fourier energy, as these areas may exhibit higher frequency patterns and more structured textures due to fluid accumulation and cellular infiltration. The negative correlation with skewness indicates that the intensity distribution in these regions is more symmetrical compared to other tissue areas. In IIA regions, tissue collapse and fluid accumulation result in a more uniform and less variable intensity distribution, leading to a reduced skewness. Additionally, the denser and more homogeneous structure results in fewer sharp boundaries or edges in the micro-CT images, which corresponds to lower LoG values and less intensity variation, resulting in lower STD values.

These results demonstrate that X-Pression accurately infers gene expression programs from micro-CT data, providing a scalable and non-destructive approach to explore ST in three dimensions.

### 3D transcriptional comparison of different vaccine strategies

In addition, we applied X-Pression to another lung specimen (L2210914) from a mouse that had been immunised with the OTS206 live-attenuated SARS-CoV-2 vaccine and challenged with the SARS-CoV-2 Delta variant for 5 days. By analysing the predicted signature composition of the sample and comparing it with the mRNA vaccine-immunised specimen, we uncovered substantial changes in the effectiveness of the different vaccines. First, we classified each voxel based on the gene expression program with the highest predicted value and assessed their spatial prevalence (**Fig.4a**). In the lung of the OTS206-immunised mouse, the gene expression program indicative of active viral replication (5) was substantially reduced across the whole organ. IIA signature (1), though present in both conditions, was diminished in the OTS206 sample, indicative of less severe infection and tissue damage. We compared the distribution of the predicted signatures, which further indicated deviations in compartments 1 and 5, as well as compartment 3 (**Fig.4b**). Signatures expressed by other structural parts of the tissue were found to show no appreciable difference.

**Fig. 4.**
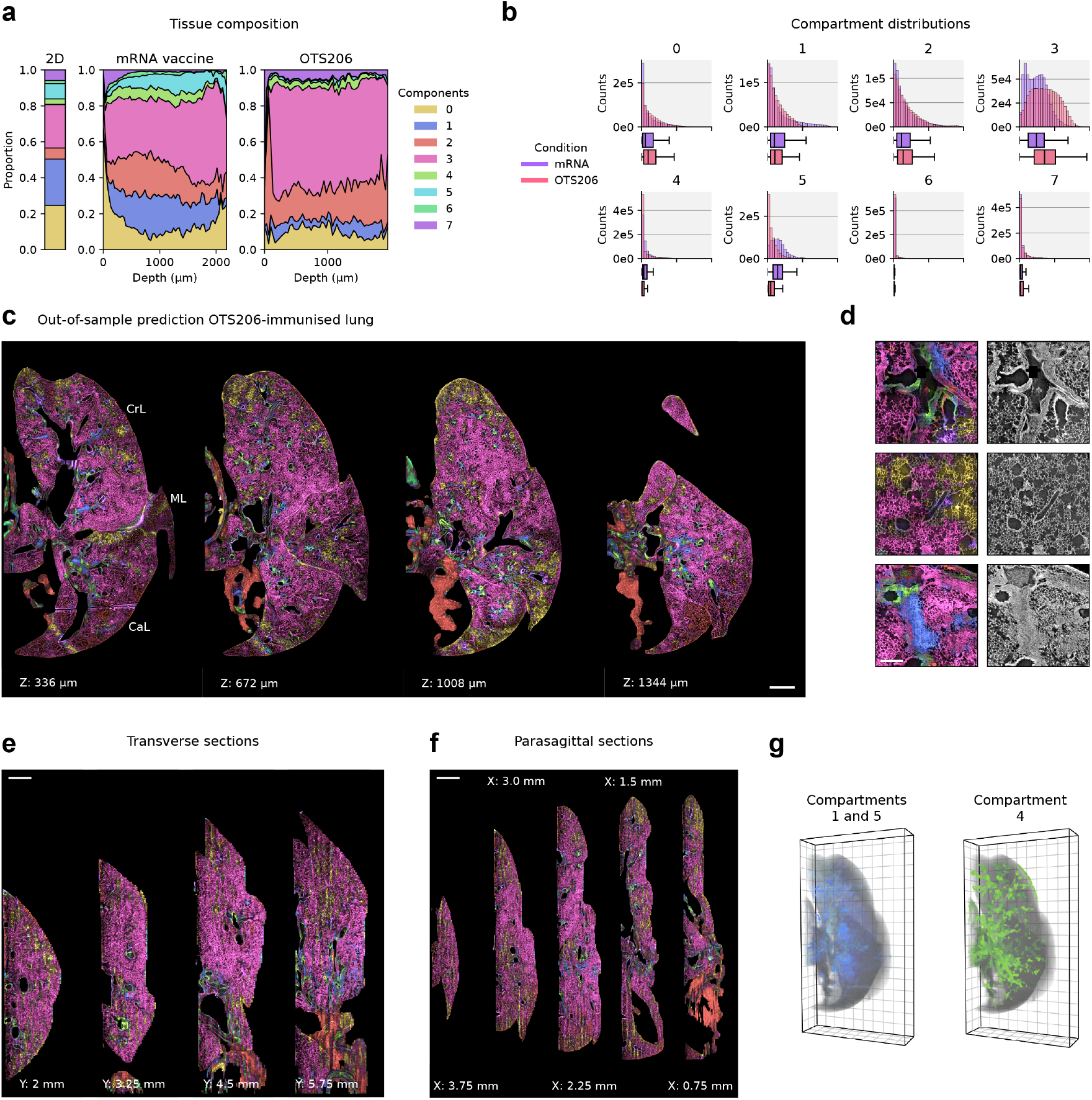
X-Pression reveals reduced viral replication and infection-associated tissue damage in a mouse lung immunised with the OTS206 live-attenuated SARS-CoV-2 vaccine. **a**, Area plots showing the proportions of the inferred 3D gene expression programs in the mouse lung immunised with mRNA vaccine or OTS206 vaccine and 28 days later challenged with the SARS-CoV-2 Delta variant. Analysed lungs were harvested 5 days post-challenge (5 dpc). Ground truth ST section (L2210926) is visualised as a stacked barplot on the left. **b**, Distribution of the predicted gene expression program scores compared across conditions, visualised as histograms and boxplots below (centre line: median, box limits: upper and lower quartiles, whiskers: 1.5x interquartile range). **c**, Serial axial sections of OTS206-immunised mouse lung with MIP colour overlays showing the gene expression signatures inferred by X-Pression. Scale bar: 1000 µm. CrL: cranial lobe, ML: medial lobe, CaL: caudal lobe. **d**, Demonstrative high-resolution ROIs with colour overlays showing the gene expression signatures and the original tomograms. Scale bar: 200 µm. **e**, Serial transverse sections with colour overlays showing gene expression changes across the volume. Scale bar: 1000 µm. **f**, Serial parasagittal sections with colour overlays showing gene expression changes across the volume. Scale bar: 1000 µm. **g**, 3D whole organ renders of expression programs associated with IIA (1) and viral replication (5) (left) and bronchioles (4) (right). Grid spacing: 500 µm.

The visualisation of the signatures further corroborated a milder infection by the SARS-CoV-2 Delta variant in the lung of the OTS206-vaccinated mouse. In this case, most of the parenchyma remains intact, while it is largely inferred to express a signature indicative of active immune infiltration (3). Although active viral replication sites were present, they were much less abundant compared to the lung of the mRNA-vaccinated animal (**Fig.4c**). Expression programs were consistent with the microstructure, showing mostly normal and moderately elevated radiodensity in the parenchyma and mild lung collapse in a few focal points (**Fig.4d**). These findings were further confirmed by visualisation of transverse and parasagittal sections (**Fig.4e-f**). 3D rendering of the whole lung with active viral replication sites and lung collapse high-lights the reduced infection severity and tissue damage compared to the mRNA-vaccinated lung (**Fig.3g**).

In summary, we show that X-Pression enables high-resolution 3D reconstruction of gene expression programs, revealing spatially distinct transcriptional landscapes within the lung tissue. By capturing complex interactions between viral replication, immune responses and tissue remodelling, X-Pression provides a powerful tool for studying gene expression dynamics across multiple scales, offering new insights into infection progression and vaccine efficacy.

## Discussion

In this study, we introduced X-Pression, a deep learning-based framework that enables scalable and cost-effective inference of 3D gene expression programs from micro-CT data. A 3D perspective is essential for a deeper understanding of tissue function and disease progression, yet most transcriptomic profiling techniques are constrained to two dimensions. While recent technological advances have significantly improved 2D ST, these two-dimensional approaches may not fully capture the complexity of three-dimensional tissue architecture, potentially leading to incomplete or inaccurate interpretations. Additionally, conventional sectioning of tissues inherently results in the loss of useful spatial information, limiting our ability to study tissue architecture and cellular interactions in their native 3D context.

To address these limitations, X-Pression presents a solution by offering high-resolution transcriptomic profiling of 3D tissue structures through an integrated approach that combines ST and micro-CT. Our method leverages deep learning-based modelling with micro-CT imaging to overcome the challenges of traditional 2D methods, resulting in a scalable and cost-effective approach. Furthermore, the non-destructive nature of micro-CT enables subsequent, targeted investigation of the tissue sample based on this predictive modelling. In addition, X-Pression allows for whole-organ analysis, thus permitting the study of gene expression patterns and a more detailed understanding of disease progression. In summary, our results demonstrate that X-Pression successfully captures information relevant to predicting gene expression from microtomograms with high accuracy and minimal data requirements from a single ST section.

Integrating higher-quality and more diverse training data will further improve the generalizability of our model, resulting in more accurate and robust predictions across a broad range of tissue types. To improve the performance of our model, we propose performing micro-CT imaging prior to ST, which would enhance the registration accuracy between H&E and CT slices, therefore improving the accuracy of gene expression predictions. Furthermore, integrating advanced model architectures, such as vision transformers (e.g., Eff-CTNet [33], Swin transformer [34]), may boost the model’s ability to generalise across different tissues. These architectures utilise attention mechanisms to capture long-range dependencies and multi-scale features, allowing the model to incorporate the global context rather than relying solely on extracted 3D patch patterns. Another promising approach would be to implement local self-attention mechanisms to improve the learning of weak textures, as demonstrated in SwinMatcher [35]. Expanding beyond micro-CT, X-Pression could also be applied to other non-destructive imaging techniques, such as regular light sheet microscopy. While these methods are less powerful than micro-CT, they remain more accessible and still provide valuable insights into tissue structure and pathology. This versatility increases X-Pression’s potential for application in diverse research settings.

To showcase the effectiveness of X-Pression, it was applied to a cohort of SARS-CoV-2-infected mice, revealing complex host-pathogen interactions. Although traditional 2D imaging and transcriptomic approaches have provided valuable insights [36], they fall short of capturing the full complexity of 3D tissue architecture and the spatial dynamics of viral replication and immune responses. Our integrated approach allowed for a detailed characterisation of viral replication sites, immune cell infiltration, and lung tissue damage at the cellular scale and provided new insights into the protective effects of different vaccine strategies.

In particular, we used X-Pression to assess how an mRNA and a live-attenuated vaccine modulate immune responses during a SARS-CoV-2 challenge infection. By analysing the spatial distribution of immune cell markers and viral replication sites, we observed distinct patterns of immune activation and protection in vaccinated animals. Our data suggest that both vaccine strategies, while differing in their mechanisms of action, are capable of promoting localised immune responses that limit viral replication in the lungs and mitigate tissue damage. The spatial separation between replication sites and collapsed regions due to inflammation suggests that immune response kinetics, rather than direct viral damage, are shaping lung pathology. Our approach highlighted that lesions (increased IIA percentage) in the lungs of a mouse previously vaccinated with an mRNA vaccine and subsequently challenged with the SARS-CoV-2 Delta variant were elevated in comparison to those in the lungs of an animal vaccinated with OTS206. The more frequent detection of these IIA lesions suggests that the live-attenuated vaccination was slightly better in preventing virus spread or effectively controlling the infection. Despite the observed differences, it is important to highlight that both the mRNA and OTS206 vaccine strategies demonstrated efficacy in activating a genetic program in the lungs that controlled the infection with the challenge of the SARS-CoV-2 Delta variant. This integrated approach provided a deeper understanding of how immune cells are mobilised within lung tissue, how vaccines influence these dynamics, and how viral infection disrupts tissue homeostasis.

By mapping immune responses and viral dynamics at an unprecedented resolution, our study highlights the potential of this framework to reveal novel therapeutic targets and inform future vaccine development. We believe that X-Pression is a powerful tool for uncovering gene expression of tissues in 3D, encompassing viral infections, cancer, and neurodegenerative diseases, among others.

## Methods

### Experimental work

#### Biosafety statement

All experiments with SARS-CoV-2 and attenuated OTS viruses were performed in biosafety level 3 (BSL3) containment laboratories at the Institute of Virology and Immunology in Mittelhäusern, Switzerland. The standard operating procedures of the BSL3 facilities have been approved by the competent local authorities.

#### Mouse experiments

A well-characterised model of SARS-CoV-2 was utilised, specifically, hACE2-K18Tg mice (Tg(K18-hACE2)2Prlmn), which were bred in the specific-pathogen-free facility of the Institute of Virology and Immunology in Mittelhäusern, Switzerland. For infection, 7-16-week-old female and male mice were anaesthetised with isoflurane and inoculated intranasally with 5,000 TCID50 (50% tissue culture infectious dose) of the SARS-CoV-2 WTD614G (BetaCoV/Germany/BavPat1/2020, Acc. No. EPI_ISL_406862). The inoculum volume was 20 µl per nostril per mouse. For vaccination experiments, mice were injected intramuscularly with a single dose of 1 µg mRNA vaccine Spikevax (Moderna) or inoculated intranasally with 5,000 TCID50 of the live-attenuated SARS-CoV-2 vaccine OTS206 [32]. Mice were challenged intranasally with the SARS-CoV-2 Delta variant (hCoV-19/Germany/BW-FR1407/2021, Acc. No. EPI_ISL_2535433) 4 weeks after vaccination. All mice were monitored daily for weight loss and clinical signs. Whole lungs were harvested for histology, micro-CT, ST, and single-cell RNA sequencing (scRNA-seq).

#### Histopathological analysis of the lungs

Histological analysis was performed on whole mouse lungs harvested at 2 and 5 days post-infection/challenge. The lungs were perfused with 4% formalin, routinary processed for histology, including embedding in paraffin, sectioned at 5 µm sections and stained with H&E.

#### Spatial transcriptomics, custom probes and gene expression analysis

Lung tissue sections of 5 µm thick were sectioned directly onto Visium Spatial for FFPE Gene Expression Mouse Transcriptome slides (6.5 × 6.5 mm) containing four capture areas each and processed according to the manufacturer’s recommendations (10X Genomics, Pleasanton, CA). In addition to the mouse transcriptome probes, we used a set of customised probes for the SARS-CoV-2 virus targeting *ORF1ab, ORF3a, ORF10* and the genes encoding the structural proteins spike (S), envelope (E), membrane (M), and nucleocapsid (N). The custom SARS-CoV-2 probes and protocol have been previously published [32]. The cDNA libraries were loaded onto the NovaSeq 6000 system (Illumina) and sequenced with a minimum of 50,000 reads per covered spot. Reads contained in Illumina FASTQ files were aligned to a custom multispecies reference transcriptome generated with Space Ranger using the GRCm38 (v.mm10-2020-A_build, 10X Genomics) mouse and NC_045512.2 SARS-CoV-2 references.

#### SR Micro-CT acquisition

We conducted synchrotron radiation-based X-ray phase contrast microtomography (SR Micro-CT) on paraffin blocks at the TOMCAT beamline (X02DA) of the Swiss Light Source (Paul Scherrer Institute, Switzerland). This imaging technique leverages the highly coherent synchrotron X-rays to detect phase shifts in a sample, enabling high-resolution, three-dimensional visualisation of internal structures with enhanced contrast. In order to image the samples completely, several microtomography scans were collected and stitched together to cover the maximum size of the sample. To localise precisely the area of interest in the paraffin block, beeswax pillars were used [37]. For the scan parameters, the TOMCAT beamline was set at a monochromatic beam energy of 21 keV, with a 100 µm Al filter and a 10 µm Fe filter. The X-ray microscope used for imaging had an effective pixel size of 1.625 µm (a PCO Edge 5.5 combined with an x4 magnification objective and a LuAG:Ce 150 µm scintillator to convert X-ray to visible light). The propagation distance used was 200 mm. The exposure time was set to 30 ms. For each scan, 3001 projections were captured, along with 30 dark images and 2×50 flat-field images. The acquisition was done using a 360° rotation of the sample to artificially increase the total field of view to 8×8×3.5 mm. The scans were completed in less than 3 minutes each. An average of 3 scans was necessary to cover one sample. Before reconstruction, the phase-retrieval algorithm [38] was used with delta = 3.7e-8 and beta = 1.7e-10 combined with an unsharp mask (stabilizer = 0.3 and width = 1). The tomographic reconstruction was then performed using the Gridrec algorithm, with output slices in an 8-bit format. The ring correction of Vo *et al*. [39] was used (Window size of 81 - 31 and SNR = 3.0).

### ST computational work

#### ST preprocessing

FastQ files were processed with Space Ranger (10x Genomics). Sequence reads were mapped to the mm10 reference transcriptome using the Mouse Transcriptome v1 probe set file. Manual fiducial frame alignment and selection of tissue-covered spots for each sample were performed using Loupe Browser (10x Genomics). We performed basic quality control (QC) on the ST data with SCANPY [40] by checking the spatial and count distributions. To filter out low-quality spots, we picked manually defined thresholds based on the overall distribution of the total counts and the number of detected genes in the capture spots. We performed normalisation using scanpy.pp.normalize_total with parameters target_sum equals to 1e4 and exclude_highly_expressed set to True. To correct for skewness in gene expression data, we applied log transformation [log(x + 1)] using scanpy.pp.log1p.

#### Histopathological annotations

Histopathological annotations were manually assessed by trained veterinary pathologists. Three distinct lung tissue regions were identified: inflammation-induced atelectasis, inflammation, and lung parenchyma. Annotation polygons were extracted from Visiopharm software (Visiopharm A/S). These polygons were mapped to ST capture spots using the Shapely Python package [41] based on whether the centroid of each capture spot fell within a given polygon.

#### Cross-talk correction between samples

During the initial stages of analysis, an issue with the indexing primers was identified, preventing the use of the i5 index for demultiplexing. As a result, demultiplexing was performed using only the i7 index (10 bp). This raised concerns about potential cross-contamination between samples, as the demultiplexing process allows for a 1bp mismatch per index by default, which could contribute to the misassignment of reads.

To evaluate the extent of cross-contamination, we focused on the expression of SARS-CoV-2-specific genes, which should be present only in infected samples. However, we detected SARS-CoV-2 marker reads in the mock-treated samples, with spatial expression patterns closely resembling those in the infected samples. This pattern suggested that these reads in the mock-treated samples were not random and likely represent contamination introduced during sample collection.

To address this cross-talk between samples, we quantified the expression levels of known SARS-CoV-2 genes (*ORF1ab, S, ORF3a, E, M, N, ORF10*) by estimating background contamination fraction from mock-treated control samples. We calculated gene contamination fractions by dividing the observed expression values by the total gene counts across all datasets. Using these values, we determined the average contamination fraction from the mock samples and computed an upper bound using a 95% confidence interval.

To eliminate contamination, we estimated per-gene contamination counts by multiplying the total gene counts by the calculated contamination fraction. We then subtracted these estimated contamination values from the expression matrix, ensuring that any resulting negative values were set to zero.

#### Gene expression program inference

We used Chrysalis to infer 8 gene expression programs from ST datasets by performing archetypal analysis on spatially variable genes (SVGs). Chrysalis identifies distinct tissue compartments by fitting a simplex to the latent representation, which is derived from the principal components (PCA) of the gene expression matrix. PCA is used to reduce the dimensionality of the gene expression data, capturing the most significant patterns of variation across genes. The simplex is then fitted to these principal components, allowing Chrysalis to find distinct tissue compartments based on key patterns of gene expression. Here, Chrysalis was trained on the 5 dpc mRNA-vaccinated sample (L2210926), instead of the whole cohort, and and the learned transformations were applied to the remaining samples. This strategy minimised the influence of signatures from other samples, preventing external noise from affecting the X-Pression’s training and performance. To infer the gene expression programs, SVGs were calculated on the normalised and logtransformed count matrix (chrysalis.detect_svgs, parameters: min_morans = 0.0, min_spots = 0.05). PCA was performed on the SVG matrix (chrysalis.pca), followed by archetypal analysis (chrysalis.aa), with the number of compartments selected based on the elbow method on the reconstruction error curve.

#### Functional characterization and downstream analysis

Pathway activity scores for the different cellular pathways and the Hallmarks MsigDB collection [42] were calculated using PROGENy [43] and decoupleR [44]. Pathway activities were inferred with a multivariate linear model (decoupler.run_mlm), using the top 500 genes ranked by pathway responses in mice. To compare pathway activity across conditions, mean pathway activities and Hallmarks were calculated by averaging the activity scores of all capture spots across each condition and visualised as heatmaps. To infer associations with cellular niches, pairwise Pearson correlation was calculated with the tissue compartment scores. Cell-type deconvolution was performed with cell2location [45]. Cell type-specific expression signatures were generated using our scRNA-seq dataset. Genes were initially filtered (adata_ref, cell_count_cutoff = 5, cell_percentage_cutoff2 = 0.03, nonz_mean_cutoff = 1.12) followed by the training of the regression model with a maximum of 1000 epochs. We trained the cell2location model (N_cells_per_location = 30, detection_alpha = 20) with epocs = 3000. Cell densities were calculated by dividing the cell type abundance values by the total number of inferred cells.

### Micro-CT computational work

#### Volume handling

We utilised Zarr [46] to store large 3D tissue image volumes in a chunked format, enabling us to store and retrieve them without loading the entire volume into memory. After extracting slices and storing them in Zarr, the 3D volumes were loaded into Dask arrays [47]. Dask was used to convert those large 3D volumes (slices from a TIFF file) into a distributed array format, allowing parallel processing to handle all data simultaneously. This approach reduced memory consumption and enhanced performance.

#### Volume segmentation

To segment the tissue from the background, we used a custom U-Net architecture implemented in Keras [48]. This architecture was trained on axial slices of tomograms, for which tissue masks were manually annotated by a pathologist and used as ground truth for supervised learning. Raw images were subsequently preprocessed by extracting 128×128 patches from the annotated slices with the corresponding binary masks. Data normalisation was performed by scaling intensity values to [0,1].

The dataset was split into training, validation, and test sets (80%/10%/10%). The U-Net model was trained using the Adam optimiser (learning_rate = 1e-4) with (1 - dice coefficient) as the loss function. Performance metrics included dice coefficient, intersection over union, recall, and precision. Training was conducted for 200 epochs with early stopping and learning rate reduction to 1e-7.

The U-Net used for this task follows an encoder-decoder structure. The encoder consists of four convolutional blocks, each containing two 3×3 convolutional layers with ReLU activation, followed by batch normalisation and 2×2 max pooling for downsampling. The number of filters in the convolutional layers increases progressively from 8 to 64. Each encoder block generates a set of feature maps that will be used as skip connections later.

At the bottleneck, two convolutional layers with 3×3 kernels, 128 filters, and ReLU activation are applied, followed by batch normalisation to prevent overfitting. The decoder mirrors the encoder structure with four upsampling blocks. Each block consists of a 2×2 transposed convolution for upsampling, followed by concatenation with the corresponding feature map from the encoder via skip connections. Skip connections are in the decoder function, where the upsampled output from the previous decoder block is concatenated with the feature map from the encoder. This allows the decoder to access high-resolution features from the encoder, helping to recover fine spatial details. Two 3×3 convolutional layers with ReLU activation and batch normalisation refine the upsampled features at each stage. The number of filters decreases symmetrically from 64 to 8.

The final output layer applies a 1×1 convolution with a sigmoid activation function to generate a segmentation mask, where pixel values represent the probability of belonging to the tissue class. The predicted patches were then stitched together to reconstruct the segmented volume. To separate tissue from the background, we applied Otsu thresholding and removed small objects from the resulting binary mask to eliminate artefacts.

#### Multimodal coregistration

To infer gene expression profiles, we coregistered the micro-CT volumes with the ST data using BigWarp [49], an interactive tool in Fiji/ImageJ for landmark-based image alignment. We decided to use BigWarp as it provides real-time visualisation of transformations. First, we loaded the slice from micro-CT volume and the corresponding H&E image from ST data into BigWarp. The micro-CT volume was set as the fixed reference image and the H&E image as the moving image. We manually selected anatomical landmarks (from 20 to 30 landmarks) that were clearly visible in both images, ensuring an accurate mapping.

To compute the spatial transformation, we extracted the landmark coordinates, used a least squares affine transformation, and applied it to the original ST spatial coordinates to align them with the micro-CT volume.

#### Convolutional neural network

We built a custom 3D CNN to predict expression programs using 3D patches extracted from the preprocessed and coregistered volumes. Our CNN architecture consists of several layers, starting with 3D convolutional layers for feature extraction. These layers use increasing filter sizes of 32, 64, 128, and 256, followed by ReLU activation. After convolution, max pooling layers were applied to downsample the feature maps. Then, the model flattens the output and passes it through a dense layer with 64 units and ReLU activation, followed by a dropout layer with a rate of 0.3 to prevent overfitting. The final output layer consists of 8 units, with a softmax activation function for multi-output regression. For model training, we used the Kullback-Leibler divergence (kl_divergence) as the loss function and the Mean Absolute Error (MAE) as the evaluation metric. The model was optimised using the Adam optimiser with a learning rate of 1e-4. Early stopping was implemented to prevent overfitting, and training was conducted for 75 epochs with a batch size of 32.

For multi-output regression, we utilised eight gene expression programs inferred by Chrysalis from the corresponding ST data. We conducted a stratified split of the dataset to address the underrepresentation of certain gene expression programs by defining stratification bins based on the highest expression program value per spot. This ensured that both the training and validation datasets had a balanced representation. The data was split into 80% for training and 20% for validation and testing. The validation and test sets were further divided into 70% for validation and 30% for testing. Augmentation techniques, including 90-degree, 180-degree, and 270-degree rotations, horizontal and vertical flips, as well as combinations of horizontal flips with each rotation, were applied to improve the model’s generalisation ability. Augmented samples were removed from the validation and test sets. The model was trained with 128×128×21 patches centred around the spatial capture spots after registration. R^2^ scores were computed to assess the model’s performance on the validation, test, and training datasets.

During inference, we extracted 3D patches from the entire volumes using a stride of 16 to predict the gene expression profiles. We reconstructed the predictions as a matrix, where each element is of the size of 16×16 pixels (26×26 µm), corresponding to the central part of each capture spot, to minimise the effect of transcript diffusion.

#### Compartment visualisation for volumetric data

To visualise the tissue compartments, we used MIP. We preselected slices from the micro-CT volume at different depths (coronal, sagittal, and axial). Then, for each slice, MIP was applied to the individual compartments and summed together with the corresponding grayscale tomogram slice to visualise the predicted gene expression programs.

To visualise each compartment separately in 3D across the entire tissue volume, we used the Imaris Viewer software by Oxford Instruments.

#### Correlation of compartments with micro-CT features

We calculated Pearson correlation coefficients between morphological features extracted from the micro-CT per patch and the predicted tissue compartments. The morphological features included texture-related measurements (skewness, dissimilarity, correlation, LoG) and intensity-related measurements (edge density, contrast, homogeneity, Angular Second Moment (ASM), energy, elongation, mean intensity, Fourier energy, STD intensity, wavelet energy). Pearson correlation coefficients were computed using the pearsonr function from scipy.stats. A correlation matrix was generated, where each entry represented the Pearson correlation between a specific morphological feature and a compartment class.

#### Integrated gradients

We applied the IG method to attribute model predictions to input features derived from a 3D patch extracted from the micro-CT image. A 3D patch centred around the X and Y coordinates was extracted and passed through the pre-trained model, and IG attributions were calculated to identify which regions of the image contributed most to the prediction. To achieve this, we first created a baseline input 3D patch with all pixel values set to zero. The input patch was then interpolated between the baseline and the actual image using alpha values ranging from 0 to 1, spaced in 50 steps. For each interpolated image, the gradients of the model’s prediction with respect to the input image were computed. Given that the model’s output is a softmax prediction, the gradients were calculated based on the argmax of the softmax output. The gradients were then averaged between consecutive pairs of interpolated images. Finally, the averaged gradients were scaled by the difference between the input image and the baseline, yielding the importance of each pixel in the image with respect to the model’s prediction. We applied PCA on the resulting attributions and performed UMAP on the first 70 PCs (59% explained variance) to visualise the structure of attribution patterns in a low-dimensional space. For UMAP visualisation, we plotted the corresponding 2D patches by extracting the middle slice from each 3D patch used in the analysis.

#### Out-of-sample predictions

For the out-of-sample predictions, we used a micro-CT volume (OTS206, 5dpc) that the model had not seen during training. To minimise inconsistencies in contrast across various scans, we applied histogram matching, using the slice employed for model training as a reference slice. For each slice in the micro-CT volume, we adjusted the pixel intensity distribution using skimage.exposure.match_histograms to match that of the reference slice.

### ScRNA-seq computational work

#### ScRNA-seq preprocessing

Raw data alignment of FASTQ files and UMIs counting were performed using Cell Ranger (10x Genomics). SCANPY was used to analyse the count matrices. Cells were marked as outliers following the single cell best practices by using rather lenient thresholds to avoid excluding smaller subpopulations, filtering cells out being on the left side (<) or the right side (>) of the median, median absolute deviations (MAD). We marked cells as outliers for log1p_total_counts, log1p_n_genes_by_counts, pct_counts_in_top_20_genes, MAD < 3 or > 5, and cells with a percentage of mitochondrial genes > 5 and MAD < 5 or > 5, and for cells with a percentage of haemoglobin genes > 3 and MAD < 3 or > 3 [50]. Following this, each sample was individually reassessed and processed for removal of doublets using scrublet [51], which simulates doublets from the cells in each sample and automatically establishes a threshold based on a k-nearest-neighbours classifier that determines which cells are estimated as doublets. PCA was applied to each sample to visualise the scores and results of the filtering steps before integration.

#### ScRNA-seq integration, clustering, and annotation

scVI [52] was used to integrate and batch-correct the data based on the batch covariate (sample) with standard parameters besides, the n_latent = 30, n_layers = 2, batch_size = 64, and n_hidden = 64 arguments, for obtaining the embeddings. These results served as input for UMAP, and clustering was conducted at various resolutions to analyse the granularity of the biological populations using the Leiden algorithm, along with the ranking of highly variable genes in each cluster at different stages. For the first round of automatic annotations, we used scimilarity [53] followed by a second round of annotations using celltypist [54] to impute the best-scoring cell types. The assigned labels were subsequently revised manually.

### Statistics and reproducibility

Statistical methods for each analysis are detailed in their respective sections. Pearson correlation coefficients were computed with the pearsonr function from SciPy’s stats module. Visualisations, including scatterplots, box plots, bar plots, stack plots, histograms, KDE plots, heatmaps, and line plots, were created using matplotlib [55], SCANPY, chrysalis, and seaborn[56]. We utilised the sklearn.metrics.r2_score function to calculate the R2 metric. For the CNN model, KL divergence was computed using TensorFlow’s reduce_mean, reduce_sum, and math.log functions. The investigators were blinded to treatment status during the experiments.

## Data availability

Raw and processed RNA sequencing data generated by Cell Ranger and Space Ranger will be available on the Gene Expression Omnibus (GEO) database in the next version of the manuscript. All processed single-cell, ST, and micro-CT data necessary to replicate the analysis presented in this work are available at the Zenodo data archive (https://doi.org/10.5281/zenodo.15064778). The uploaded dataset includes count matrices, AnnData objects, as well as results for cell type deconvolution, tissue compartment inference, and additional metadata.

## Code availability

X-Pression is available at https://github.com/rockdeme/x-pression.

## Acknowledgements

We thank H.Y. Stoller-Kwan from the Institute of Hospital Pharmacy at the University Hospital of Bern and U. Romanelli from the Department of Infectious Diseases, Tropical Medicine and Travel Medicine at the University Hospital of Bern for providing the mRNA vaccine. We thank the Next Generation Sequencing Platform of the University of Bern for performing the high-throughput sequencing experiments and the COMPATH platform of the Institute of Pathology and the Institute of Animal Pathology of the University of Bern for performing the pathological analyses. We acknowledge the Paul Scherrer Institut, Villigen, Switzerland for the provision of synchrotron radiation beamtime at the TOMCAT beamline X02DA of the SLS. This project was supported by the Novartis Foundation for Medical-Biological Research (grant 22B099 to E.A.M.).

## Author Information

**Contributions.** D.T., L.G., V.T. and E.A.M. designed the study. D.T. and L.G. designed computational models, algorithms, and validation. E.A.M. developed the experimental methodology and validation. E.A.M., N.E. and G.T.B. performed the mice experiments. M.B. performed scRNA-seq analysis. I.B.V., L.GR. and C.H. performed histopathological validation and tissue annotations. A.B. acquired the micro-CT images. A.C. performed tissue sectioning and ST experiments. D.T., L.G. and E.A.M. drafted the manuscript. A.V., S.R., V.T., and E.A.M. supervised the research. All authors revised and approved the published version of the manuscript.

## Ethics declarations

Mouse studies were approved by the Animal Experiments Committee of the Cantonal Veterinary Office of Bern and conducted in accordance with Swiss animal welfare legislation under licence BE43/20.

