## Supplementary Information for "Deep learning-based 3D spatial transcriptomics with X-Pression"

### Supplementary Note 1: characterisation of gene expression signatures

To characterise the gene expression programs (*i.e.*, molecular tissue compartments) presented in our study, we relied on directly examining the highest contributing genes to each compartment, performed cell type deconvolution using our single-cell RNA sequencing (scRNA-seq) reference, and histopathological assessment.

Compartment 0 was mainly associated with alveolar epithelium. *Sftpc* (Surfactant protein C), *Ager* (Advanced glycation end-product receptor), and *Tmem100* (Transmembrane protein 100) are markers for alveolar type I and II cells. *Hbb-bs*, *Hbb-bt*, and *Hba-a2* haemoglobin genes indicated the presence of erythroid cells. *Epas1*, *Igfbp2*, *Cavin2*, and *Inmt* are linked to endothelial cells and vascular function.

Compartment 2 was linked to fibroblasts and complement-associated immune response, extracellular matrix remodelling, complement activation, and immune cell interactions. Fibrosis and extracellular matrix remodelling were marked by *Col1a1*, *Col1a2*, *Col3a1*, and *Fabp4*. We further observed complement genes (*C3*, *C4b*, *Cfd*, *Cfp*), tissue repair (*Igfbp4*), and chemokine genes (*Ccl6*) associated with macrophage recruitment.

Compartment 3 suggested parenchymal epithelium components and inflamed alveolar tissue. *Sftpc* and *Ager* again point to type II and type I pneumocytes. Inflammatory markers, such as *Lcn2*, *Cxcl15*, and *Saa3* pointed to inflammation and immune responses in the tissue. *Krt7*, *Spock2*, and *Apod* highlighted epithelial regeneration and remodelling.

Compartment 4 was associated with bronchiolar epithelium and club cells based on the presence of *Scgb3a2*, *Cyp2f2*, and *Wfdc2*. *Hp*, *Trf*, and *Pigr* indicate vascular components as well. Immune activation related genes can also be found, such as *Cxcl17* and *Retnla*.

Compartment 6's signature indicated contractile tissue (*Myh6*, *Myl7*, *Tnnt2*) and structural proteins (*Ankrd1*, *Tcap*, *Csrp3*).

Compartment 7 marked epithelial and smooth muscle cell-related genes. Club cell markers (*Cyp2f2*, *Cbr2*, and *Aldh1a1*), smooth muscle genes (*Acta2*, *Tagln*), and epithelial markers (*Scgb3a2*, *Wfdc2*, and *Sec14l3*) together suggested an expression program related to larger airways.

Cell type deconvolution was performed using paired scRNA-seq data we acquired at 2dpi and 2dpc for the OTS206 vs. Wuhan infection and the OTS206 vs. mRNA vaccine challenge experiments, respectively (**Supplementary Fig.1a-c**). Next, we calculated the correlation between compartment scores calculated with Chrysalis and the cell type proportions. On the 5 dpc mRNA-vaccinated sample (L2210926) used for training, we revealed that pneumocytes, endothelial cells, and other parenchymal cell types showed positive association with structural compartments, whereas immune cells, such as CD4+ T cells, CD8+ T cells, and B cells, correlated with compartments 1, 2, 3. Furthermore, the active viral replication signature (5) correlated with neutrophils and macrophages.

Moreover, we assessed the compartment composition of annotated capture spots across the whole ST dataset (**Supplementary Fig.1d**). This showed that spots annotated as parenchyma contained the highest fraction of scores for compartments 0, IIA for compartment 1, inflammation and prominent inflammation for compartment 5, and other structural components in the unlabelled category for the rest of the compartments.

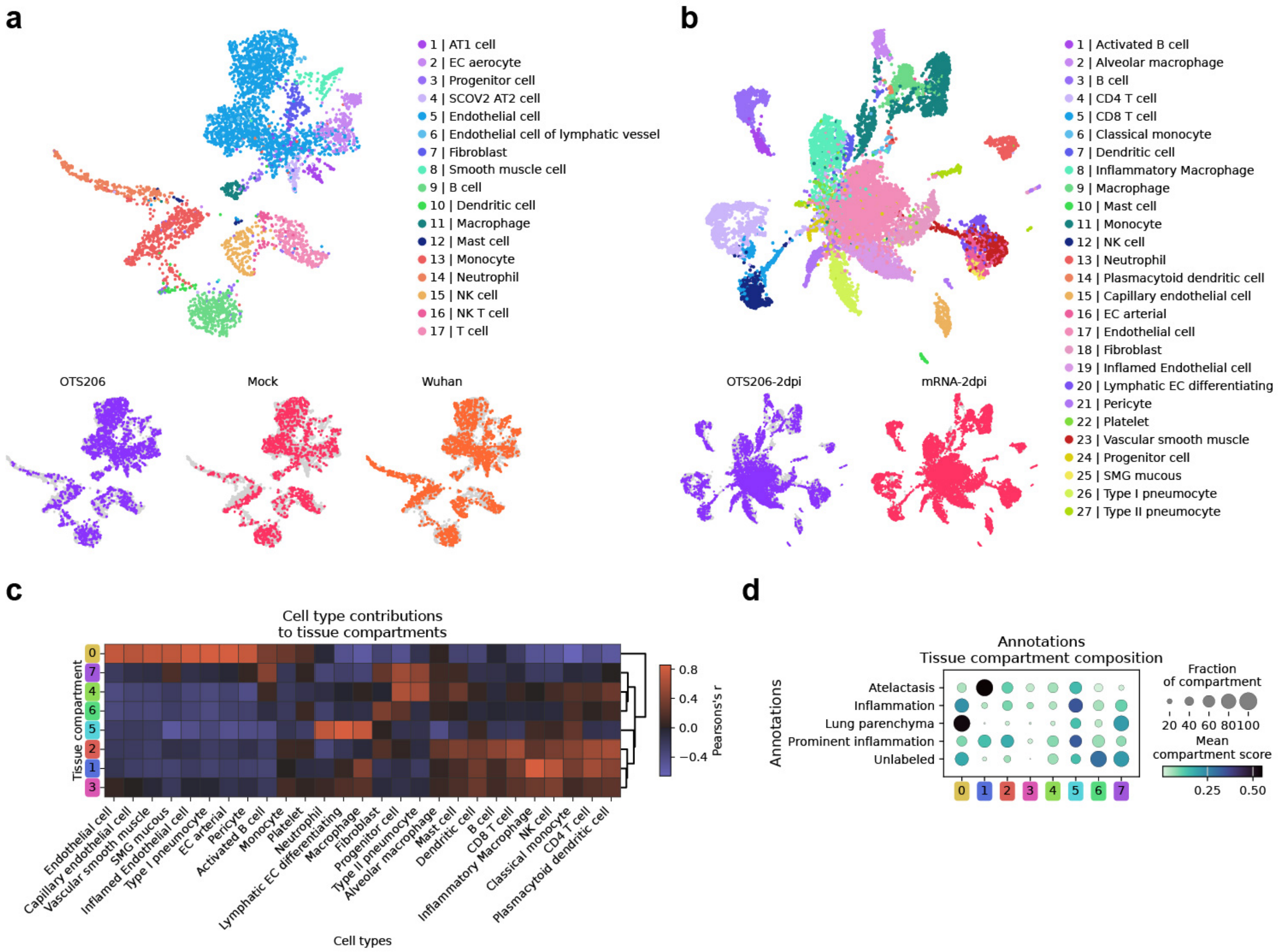


**Supplementary Fig. 1. | Characterisation of gene expression programs through cell type deconvolution and histopathological annotations. a,** Uniform manifold approximation and projection (UMAP) representation of single cells collected from mouse lung tissues two days after infection of the Wuhan SARS-CoV-2 strain and OTS206. **b,** UMAP representation of single cells collected from mouse lung tissues two days after OTS206 and mRNA-vaccinated animals were challenged with the Delta variant. **c,** Heatmap displaying the correlation between cell type and tissue compartments in the L2210926 sample. **d,** Dotplot showing the tissue compartment composition of each annotated region in the L2210926 sample.

### Supplementary Note 2: spatiotemporal changes in the transcriptional landscape of SARS-CoV-2

To assess gene expression changes in the ST cohorts, we examined the gene expression programs inferred with Chrysalis. In the first cohort of mice infected with OTS206 or the Wuhan strain, tissue samples were taken at 2 dpi and 5 dpi (**Supplementary Fig.2a**). By evaluating the spatial distribution of IIA and viral replication-associated compartments, we observed that active viral replication was significantly more prominent in the animals infected with the Wuhan strain, at both 2 and 5 dpi, compared to those infected with OTS206 (**Supplementary Fig.2b**). The IIA-associated signature was also slightly increased in the Wuhan strain-infected animals (**Supplementary Fig.2c**). These changes were further confirmed by examining the cumulative fractional values of these compartments (**Supplementary Fig.2d**). Furthermore, we assessed the expression changes in the second cohort of mice, which were vaccinated with either the OTS206 or an mRNA vaccine 28 days prior to being challenged by the Delta variant. Samples were taken at 2 dpc and 5 dpc (**Supplementary Fig.2e**). Here, we noted that the IIA-associated signature was more pronounced in the mRNA-vaccinated animals, particularly at 5 dpc (**Supplementary Fig.2f**). Examination of the viral replication compartment revealed a more significant presence of this signature in the mRNA-vaccinated animals, especially at 2 dpc, compared to the OTS206-immunised animals (**Supplementary Fig.2g-h**).

Given that we observed substantial transcriptional changes, we extended our analysis by assessing pathway activities and gene set environments. First, pathway activity scores were inferred using PROGENy for each capture spot. We then calculated the correlation between these activity scores and the gene expression programs. This revealed a strong association of the active viral replication compartment with the JAK-STAT, NFkB, and PI3K pathways in both cohorts (**Supplementary Fig.3a-b**). Furthermore, we observed similar trends with the IIA signature, but only in the challenge cohort. By assessing the mean pathway activity scores across conditions, we uncovered a significant temporal change in JAK-STAT pathway activity. In the infection cohort, animals infected with the Wuhan strain exhibited upregulated JAK-STAT activity as early as 2 dpi, which remained prominent at 5 dpi, whereas the OTS206-infected animals showed comparable values only at 5 dpi (Supplementary Fig.3c). In the challenge cohort, both immunisation strategies resulted in similar increases in JAK-STAT activity at 2 dpc, which diminished by 5 dpc (**Supplementary Fig.3d**).

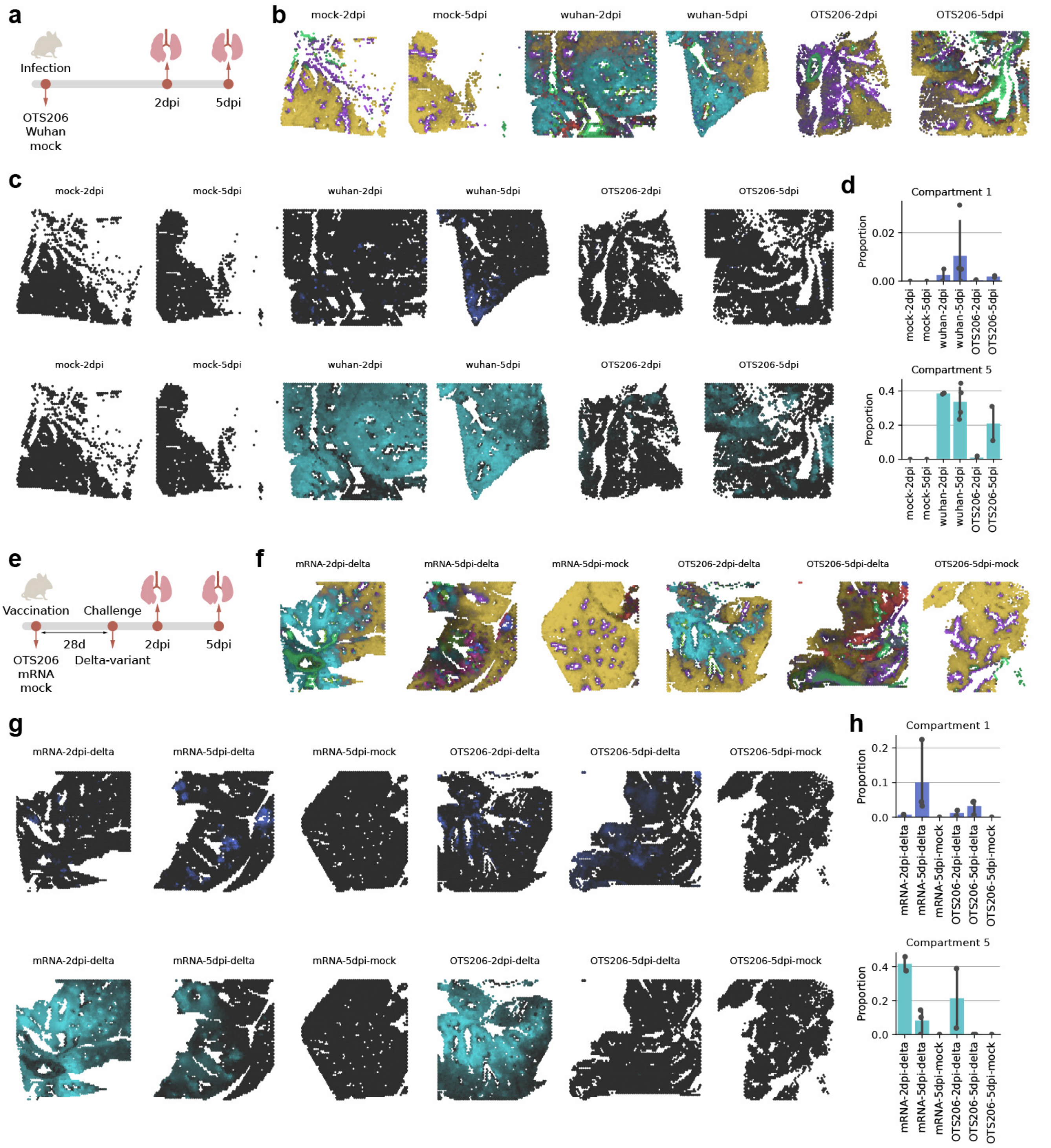


**Supplementary Fig. 2. | Spatial and temporal variations in the transcriptional landscape of SARS-CoV-2. a,** Animals were infected with the SARS-CoV-2 Wuhan strain or with OTS206. Lung tissues were harvested two and five days post-infection. **b,** MIP of the identified tissue compartments on representative tissue sections from the infection cohort. **c,** Spatial plots of compartment scores of IIA (1) and active viral replication (5) on representative tissue sections from the infection cohort. **d,** Proportions of IIA (1) and active viral replication (5) compartments across all tissue samples from the infection cohort (bar height: mean compartment fraction, black dots: individual ST samples, error bar: 95% CI, bar height: mean value). **e,** Animals were first immunised with OTS206 or an mRNA vaccine, followed by the infection of the SARS-CoV-2 Delta variant after 28 days (challenge). Lung tissues were harvested two and five days post-challenge. **f,** MIP of the identified tissue compartments on representative tissue sections from the challenge cohort. **g,** Spatial plots of compartment scores of IIA (1) and active viral replication (5) on representative tissue sections from the challenge cohort. **h,** Proportions of IIA (1) and active viral replication (5) compartments across all tissue samples from the challenge cohort (bar height: mean compartment fraction, black dots: individual ST samples, error bar: 95% CI, bar height: mean value).

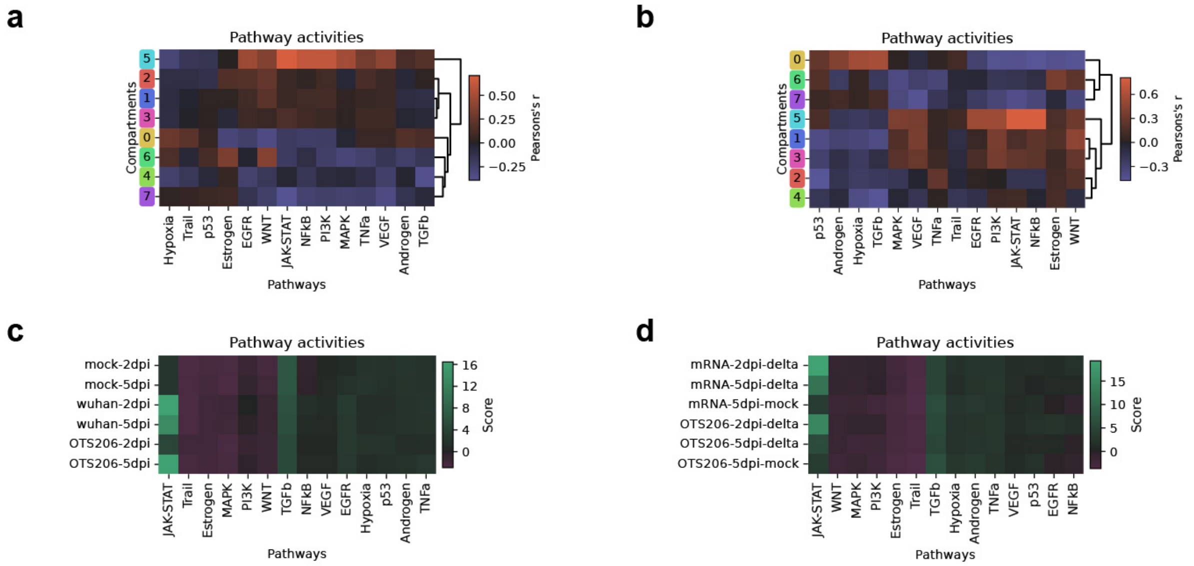


**Supplementary Fig. 3.** | **Pathway activities inferred with PROGENy. a,** Heatmap displaying the correlation of tissue compartments with intracellular pathway activities in the infection cohort. **b,** Heatmap displaying the correlation of tissue compartments with intracellular pathway activities in the challenge cohort. **c,** Heatmap depicting changes in intracellular pathway activities across various time points in the infection cohort. **d,** Heatmap depicting changes in intracellular pathway activities across various time points in the challenge cohort.
